# Designed Anchoring Geometries Determine Lifetimes of Biotin-Streptavidin Bonds under Constant Load and Enable Ultra-Stable Coupling

**DOI:** 10.1101/2020.05.12.090639

**Authors:** Sophia Gruber, Achim Löf, Steffen M. Sedlak, Martin Benoit, Hermann E. Gaub, Jan Lipfert

## Abstract

The small molecule biotin and the homotetrameric protein streptavidin (SA) form a stable and robust complex that plays a pivotal role in many biotechnological and medical applications. In particular, the biotin-streptavidin linkage is frequently used in single molecule force spectroscopy (SMFS) experiments. Recent data suggest that biotin-streptavidin bonds show strong directional dependence and a broad range of multi-exponential lifetimes under load. Here, we investigate engineered SA variants with different valencies and a unique tethering point under constant forces using a magnetic tweezer assay. We observed two orders-of-magnitude differences in the lifetimes, which we attribute to the distinct force loading geometries in the different SA variants. We identified an especially long-lived tethering geometry that will facilitate ultra-stable SMFS experiments and pave the way for new biotechnological applications.

## Introduction

The non-covalent, high-affinity binding of the small molecule biotin to streptavidin (SA) is ubiquitously used in a variety of biological, chemical, biophysical and pharmaceutical applications^1, 2, 3^. Biotin can readily be covalently attached to nucleic acids^4, 5, 6^, proteins^7, 8^, or linker molecules^9^. SA is stable over a wide range of conditions and easy to handle^1^. Owing to the specificity of the binding, as well as the robustness of the complex, the interaction has in particular become a popular tool in the context of single-molecule force spectroscopy (SMFS) assays^10^. It serves as a molecular handle to anchor molecules of interest and apply forces and torques to them^5, 11-18^. The long lifetime of the SA-biotin complex under external forces has enabled constant-force SMFS experiments lasting for hours and even up to weeks in magnetic tweezers (MT)^19, 20^. Despite its widespread use, SA’s tetravalency poses a problem, in particular in SMFS applications, since it is *a priori* ambiguous which of the four subunits biotin binds to. This ambiguity results in four different force-loading geometries for a given attachment of the SA tetramer (**Figure 1**)^21^. Furthermore, if SA is non-specifically attached − as is the case in many commercially available SA-coated magnetic beads − a variety of attachment points combined with tetravalency results in an even larger range of possible force-loading geometries^20, 22^.

**Figure 1.**
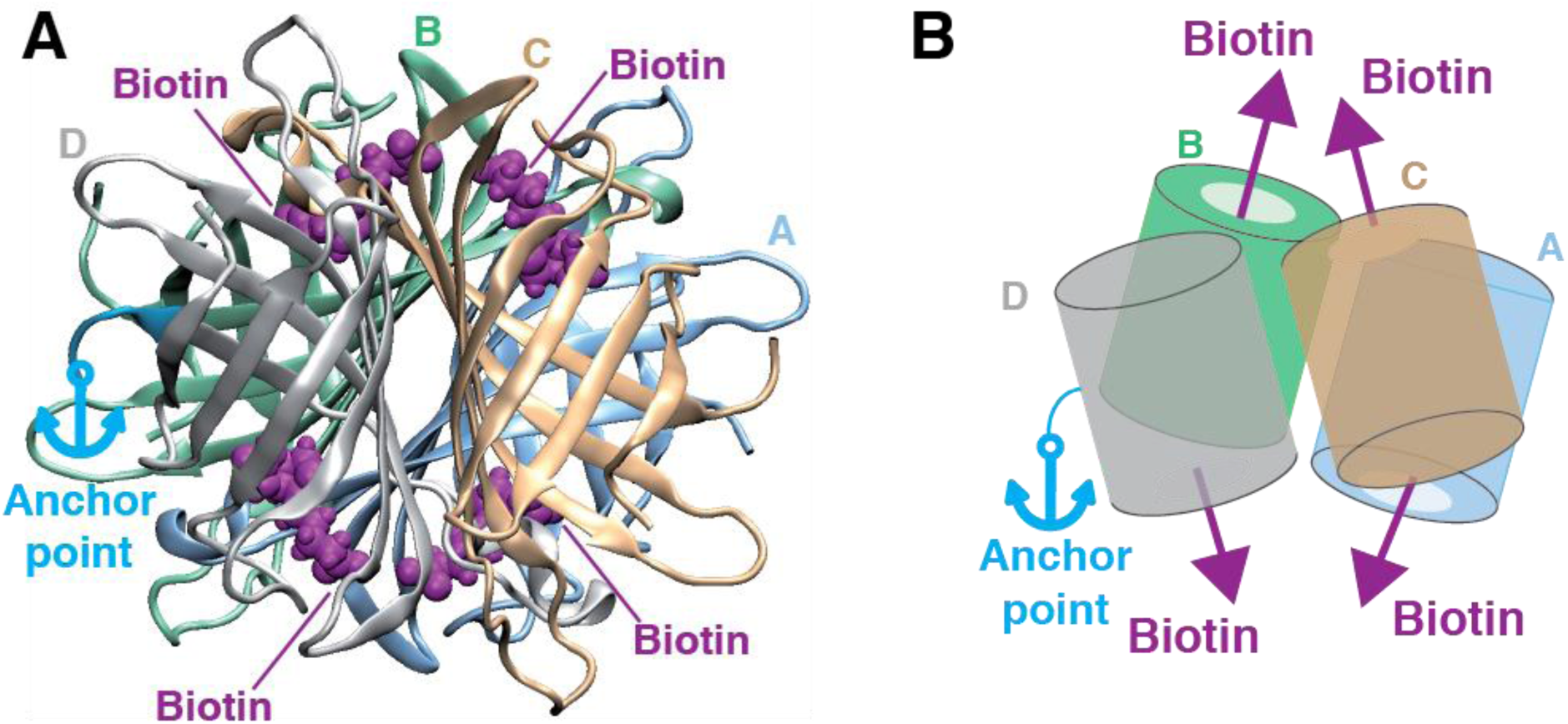
SA’s tetravalency results in different force-loading geometries. **A** crystal structure of the SA tetramer (PDB-ID: 6M9B^36^, rendered using VMD^37^) with the four subunits shown in different colors. Four bound biotin molecules are shown in purple. The light blue arrow marks the anchor point (C-terminus of subunit D). **B** Schematic representation of the tetramer structure. The colored barrels represent the four subunits. Arrows indicate the initial force-loading directions in the SMFS experiments: The light blue arrow marks the C-terminus of subunit D used for site-specific immobilization. Purple arrows indicate the four possible directions of pulling biotin out of the different binding pockets. Under constant load, the complex will rotate and rearrange in such away that the sum of forces acting on it equals zero. Depending on which subunit biotin is bound to, the orientation of the complex will be different resulting in different force propagation pathways.

Atomic force microscopy (AFM)-based constant speed SMFS experiments have recently shown that the force needed to unbind biotin from the SA binding pocket is strongly dependent on the force-loading direction^23, 24^: Tethering SA by a single defined residue and pulling biotin out of one of the binding pockets results in different force-loading geometries, depending on which SA subunit the biotin has bound to. For some of the pulling directions, the SA subunit is deformed such that the energy barrier of the binding is decreased, causing lower biotin unbinding forces^24^. However, the influence of the tethering geometry of SA on the lifetime of the SA-biotin interaction under constant forces is currently unknown.

Here, we employ engineered variants of SA with different defined valencies and a unique tethering point to restrict and control the number of possible force-loading geometries for SMFS measurements. We use AFM imaging to verify the valencies by showing that only the competent subunits can bind biotin. Furthermore, we employ isothermal titration calorimetry (ITC) to directly measure the binding enthalpies of the different SA variants. With an MT assay we assess the stability of the SA-biotin interaction under constant load and demonstrate large differences in the lifetime depending on the attachment geometry. The different stabilities give rise to multi-exponential lifetime distributions for multivalent constructs. By using one well-defined geometry and a monovalent SA construct, a single-exponential lifetime is achieved. We expect our results to be highly relevant for force spectroscopy, and, in general, to improve assays where the SA-biotin bond is under load, *e.g*. through fluid flow or rinsing steps.

## Results and Discussion

To systematically investigate the stability of the SA-biotin complex under constant mechanical load, we prepared tetra-, tri-, and monovalent variants of SA. These comprise four, three, and one functional subunit(s), while the remaining subunits are incapable of biotin binding (4SA, 3SA, and 1SA; **Figure 2A**) due to three mutations located around the binding pocket (N23A, S27D, S45A)^25^. In addition, a variant consisting of four non-functional subunits (0SA) was prepared. All variants possess a single cysteine residue at the C-terminus of their subunit D, allowing for site-specific immobilization^14, 21, 24^. For 3SA and 0SA, subunit D is non-functional, whereas for 1SA and 4SA, it is functional (**Figure 2A**; for details on protein engineering see Supplementary Materials and Methods).

**Figure 2.**
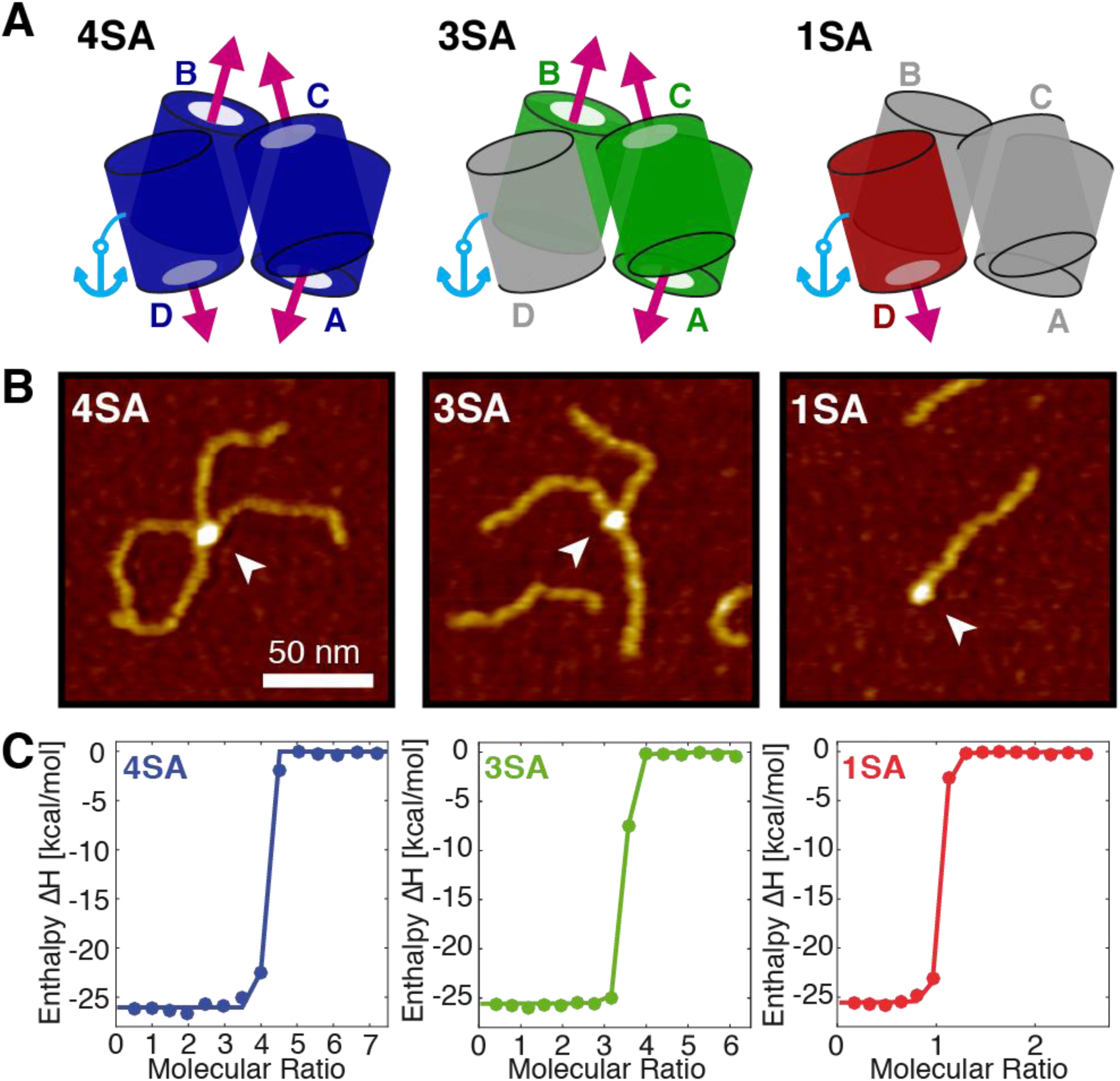
Probing SA variants with different valencies by AFM and ITC. **A** Schematic structure of SA constructs with different valencies. 4SA (left), 3SA (middle), and 1SA (right) have four, three and one functional subunit(s) (colored), respectively. The remaining subunits (gray) are incapable of binding biotin. All constructs have a single C-terminal cysteine at their subunit D –nonfunctional for 3SA, functional for 1SA and 4SA– for site-specific immobilization (light blue line). The light blue anchors mark the anchoring site of SA for the SMFS experiments, while the purple arrows indicate the possible directions in which biotin can be pulled out of the binding pockets. **B** AFM images of 4SA (left), 3SA (middle), and 1SA (right) with the maximal number (four, three, and one, respectively) of biotinylated DNA strands bound. Arrows mark the SA molecules. Height range of color scale is 2 nm. **C** Isothermal titration calorimetry data of free biotin binding to SA of different valencies. Colored dots are the measured heat release per mole upon adding biotin to SA plotted against the molecular ratio (biotin per SA) in the measurement cell. Lines are fits to the data (taking the discreteness of the measurement into account). For details of the fits see Supplement.

### AFM imaging reveals binding stoichiometry

To verify the valency of the different variants, we incubated them with biotinylated 250 bp double-stranded DNA constructs and directly visualized the resulting SA-biotinylated DNA complexes by AFM imaging (**Figure 2B** and Supplementary **Figures S1-S4**). An excess of biotinylated DNA over SA (approximately twenty-fold for 4SA and 3SA, and four-fold for 1SA and 0SA) was used to ensure that SA molecules with DNA strands bound to all functional subunits could be observed. Indeed, a maximum of four, three, and one bound biotinylated DNA strand(s) was observed for 4SA, 3SA and 1SA, respectively, confirming the expected valencies (Supplementary **Figure S5**). In the case of 0SA, no SA-biotinylated DNA complexes were observed.

### Thermodynamic parameters determined by isothermal titration calorimetry

Next, we performed isothermal titration calorimetry (ITC) measurements to determine the thermodynamic parameters of binding for biotin to the different constructs in the absence of force (**Figure 2C**). In principle, ITC allows determination of the stoichiometry, the affinity, and the binding enthalpy. To ensure good comparability across measurements, we used the same biotin stock solution with an estimated 5% uncertainty in absolute concentration for all measurements. The uncertainty in the concentrations of the SA stocks was estimated to be 10% (see Supplementary Materials and Methods). Fits to the ITC data give values for the binding stoichiometries of 1.0±0.2 for 1SA, 3.3±0.5 for 3SA, and 3.9±0.6 for 4SA (**Figure 2C**), in excellent agreement with the results from AFM imaging (**Figure 2B** and Supplementary **Figure S5**). The largest contributions to the measurement errors result from the uncertainties in concentration and the uncertainties of the values increasing with the number of available binding sites, because a given uncertainty in protein concentration has a larger impact on the uncertainty with increasing stoichiometry. Due to limitations of our instrument and the very high affinity of biotin to streptavidin, the binding constant could not be obtained directly and we can only provide an upper limit of K_D_<1 nM. We obtained binding enthalpies per binding site of −(25.0±1.3) kcal/mol for 1SA, −(25.6±1.4) kcal/mol for 3SA and −(26.1±1.3) kcal/mol for 4SA (Supplementary **Figure S6**). These results agree well with enthalpies measured in previous studies^21, 26, 27^. Within experimental errors, the binding enthalpies per binding site for all SA variants are the same, suggesting that in the absence of force all subunits are equivalent with regard to biotin binding and that no effects of binding geometries or binding cooperativity come into play.

### Single-molecule MT measurements determine lifetimes under force

To directly measure the lifetimes of the SA-biotin interactions under constant force and to investigate the influence of different force-loading geometries, we performed MT measurements using the different SA variants (**Figure 3**). In MT, the molecular construct of interest is tethered between the bottom surface of a flow cell and a superparamagnetic bead (**Figure 3A**). By applying a magnetic field, generated by permanent magnets, a constant force is exerted on the bead and thereby on the tether^28, 29^. We track the 3D position of the bead and the extension of the tether can be determined with nanometer resolution. Importantly, with our MT setup we can track approximately 100 beads in parallel, enabling us to obtain good statistics in a short amount of time^20^. In addition, MT provide excellent force and drift stability, facilitating long measurements^20^, which are critical due to the high stability of the biotin-SA bonds.

**Figure 3.**
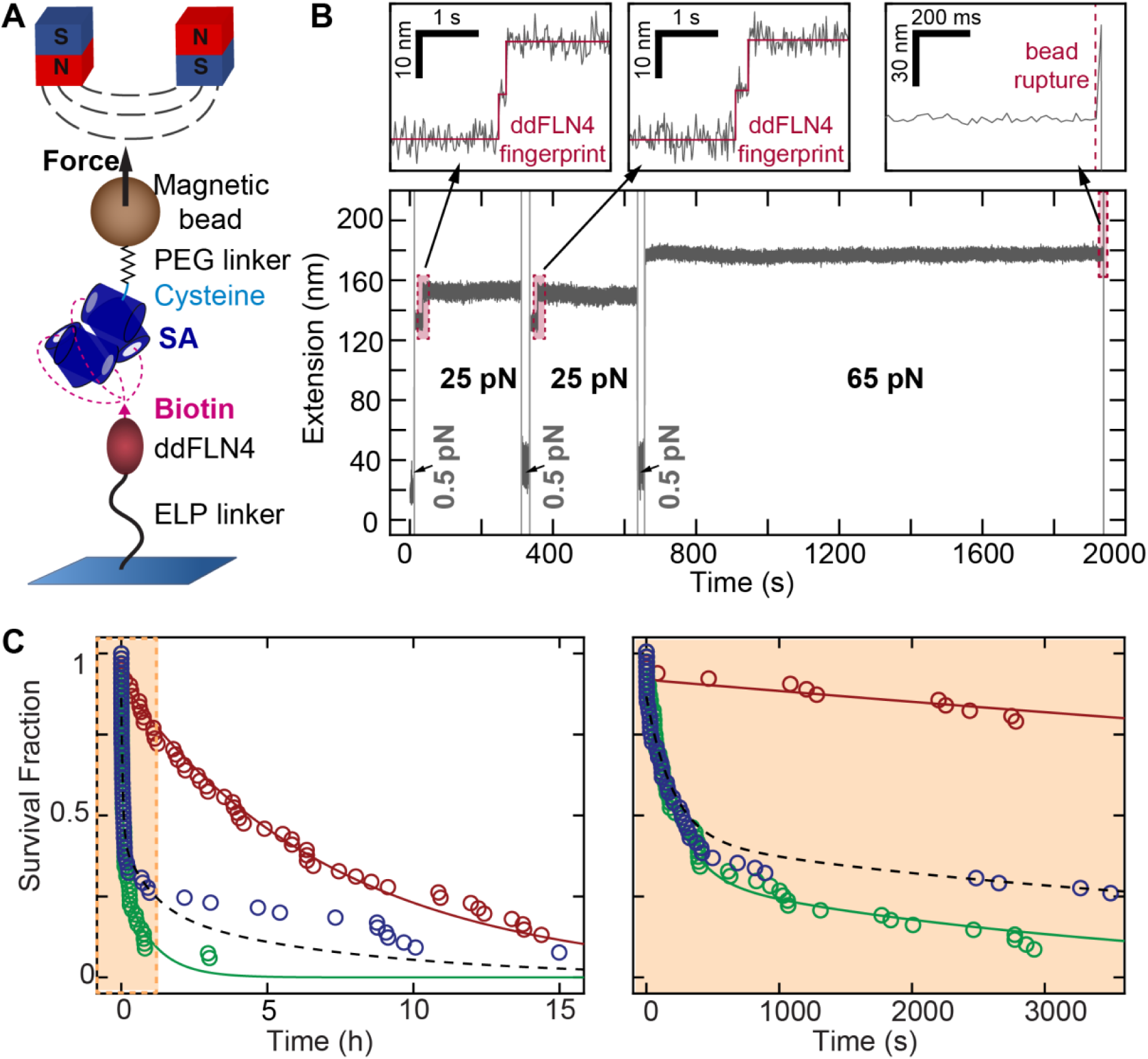
Lifetimes of SA-biotin interactions under constant force. **A** Schematic of MT experiments (not to scale). SA (4SA, 3SA, or 1SA) is site-specifically and covalently immobilized on magnetic beads via the single C-terminal cysteine at its subunit D using a PEG linker with a thiol-reactive maleimide group. Biotinylated ddFLN4 is covalently immobilized on the bottom slide of the MT flowcell via an ELP peptide linker. Binding of the biotin to one of the functional subunits of the respective streptavidin construct tethers the beads by a single SA-biotin bond. Force is exerted on the magnetic beads by permanent magnets positioned above the flowcell. **B** Time trace of the tether extension during an MT measurement. At the beginning of the measurement, beads are subjected to two 5-min intervals at 25 pN, to observe unfolding of ddFLN4 in a characteristic two-step pattern (left and middle zoom-in), which serves as fingerprint to identify specific, single-tethered beads. Short low force intervals (0.5 pN) allow for ddFLN4 refolding. Tethers are then subjected to a constant force of 65 pN and the time until bead rupture due to unbinding of biotin from streptavidin is monitored (right zoom-in). **C** Survival fractions at 65 pN as a function of time for 1SA (red), 3SA (green), and 4SA (blue). The 1SA data were fit with a single exponential (red line) with a mean lifetime of 2.6·10^4^ s. The 3SA data were fit with a two-component model (Supplementary Materials and Methods) with a short lifetime of 199 s and a long lifetime of 3.6·10^3^ s (green line). The 4SA data are well described by the predicted response from the combination of the 1SA and 3SA lifetimes that takes into account the binding site stoichiometry (black dashed line, not a fit). The left panel shows a zoom on the first hour of the data.

For the MT measurements, the small protein domain ddFLN4 (fourth F-actin cross-linker filamin rod domain of *Dictyostelium Discoideum*^30^) was biotinylated and covalently coupled to the bottom surface of a flow cell by an elastin-like polypeptide linker^31^. The different SA variants (4SA, 3SA, or 1SA) were site-specifically and covalently immobilized on magnetic beads *via* polyethylene glycol (PEG) linkers, by reacting the C-terminal cysteine of subunit D with a thiol-reactive maleimide group on the PEG linker (**Figure 3A**). The SA-functionalized beads were introduced into the flow cell and one of the functional subunits of the respective SA construct bound to the biotinylated ddFLN4, thereby tethering the magnetic bead to the surface. Upon force application, the molecular linkers are stretched and ddFLN4 unfolds in a characteristic two-step manner^20, 32^. We use the distinct two-step unfolding pattern as fingerprint to identify specific, single-tethered beads, *i.e*. beads that are bound to the surface *via* a single SA-biotin interaction. To measure the lifetime of this interaction, beads were subjected to a constant force of 65 pN and the time until bead rupture was recorded. The rupture events are attributed to the unbinding of biotin from SA, as this is the only non-covalent bond within the tether connecting beads and the surface.

Measurements of 1SA, the monovalent variant, exhibited a survival time distribution that is well described by a single-exponential fit (**Figure 3C**, red) with a lifetime of τ_1_ = 7.2 h ± 0.2 h (2.61·10^4^ s ± 680 s; see Materials and Methods for details of the fits). The fitted lifetime is in good agreement with the 6.7 h reported recently for a smaller data set^20^. The single-exponential lifetime suggests the presence of a single population, fully consistent with the expectation that for 1SA only subunit D (attached to the bead *via* its C-terminus) is capable of binding biotin. All 1SA functionalized beads are thus tethered in the same geometry, resulting in one well-defined force-loading direction.

3SA is complementary to the 1SA variant, in the sense that all but the attached subunit are functional, so that three different pulling geometries are possible for 3SA. The lifetime measurements reveal much shorter overall lifetimes for 3SA compared to 1SA (**Figure 3C**, compare red and green data points). In addition, the data are not well described by a single exponential, suggesting that the different possible pulling geometries for 3SA give rise to different lifetimes. The 3SA data are well described by a fit with the sum of three exponentials and we find two relatively short and one longer lifetime (fitted lifetimes are 98 s ± 12 s, 365 s ± 30 s, and 3100 s ± 230 s). A simpler model that combines the two shorter lifetimes into one exponential, f(t) = f_2_/3 [1 exp(−t/τ_2_) + 2 exp(−t/τ_3_)], fits the 3SA data almost equally well (**Figure 3C**, green). From the fit we obtain the two distinct lifetimes as τ_2_ = 36<u>10</u> s ± 350 s and τ_3_ = 199 s ± 10 s, where the weighting factors of the fit formula are chosen such that two thirds of the interaction of biotin and 3SA exhibit the short lifetime and one third exhibits the long one. We hypothesize that the longer-lived population corresponds to binding to one subunit, while the other two subunits exhibit lifetimes under mechanical tension that are similar.

Using steered molecular dynamics simulations, Sedlak *et al.*^24^ recently showed that for pulling biotin out of subunit A or C of SA tethered by the C-terminus of subunit D, the molecular linker adjacent to biotin gets pushed against a flexible peptide loop, significantly lowering the mechanical stability of the binding pocket. For pulling biotin out of subunit B, the same effect occurs, yet markedly less pronounced. Therefore, we assign the longer lifetime for the 3SA construct to biotin unbinding from subunit B. The shorter-lived population that comprises approximately two thirds of all unbinding events is consequently assigned to the sum of unbinding events from subunits A and C. Remarkably, the lifetime of this shortest-lived population at 65 pN is 130-fold lower than the one observed for the 1SA construct and even the long lifetime observed for 3SA is still approximately one order of magnitude shorter than the lifetime observed for 1SA.

For 4SA, we observe a rapid initial decay of bonds, but also very long-lived tethers (**Figure 3C**, blue). Since for 4SA all force-loading geometries realized in 3SA and 1SA are possible, we expect a combination of the short-lived populations observed for 3SA and the long-lived population observed for 1SA constructs. Based on this assumption, we co-plotted a prediction for the 4SA survival fraction over time as given by f(t) = f_3_/4 [1 exp(−t/τ_1_) + 1 exp(−t/τ_2_) + 2 exp(−t/τ_3_)] using the lifetimes obtained from fitting the 1SA and 3SA data (**Figure 3C**, black dashed line). The prediction using the fitted lifetimes from the 1SA and 3SA data closely matches the experimentally determined lifetimes for the 4SA variant, confirming the validity of our lifetime model and suggesting essentially random binding to the different subunits.

Combined, the above findings confirm the hypothesis that the lifetime of biotin unbinding from SA under constant force strongly depends on the tethering geometry. This finding agrees with results obtained by AFM-based constant speed SMFS experiments and can likely be attributed to the same molecular mechanism: For certain pulling directions, the SA binding pocket itself is deformed before biotin leaves the pocket. This alters the energy landscape of the binding and results in lower unbinding forces for measurements at constant retraction velocities^24^, and in shorter lifetimes for constant force experiments.

More importantly, from an application perspective, the force-loading geometry that yields the longest lifetime corresponds to pulling biotin out of the binding pocket of the subunit that is C-terminally tethered. The lifetime for this geometry is, at the force probed here, almost two orders-of-magnitude higher than for the other possible geometries. Thus, it is highly beneficial to utilize this geometry in applications for which high yield of tethers with high force stability is desirable. Importantly, this can straightforwardly be realized employing the 1SA variant used in our experiments.

Finally, it is noteworthy that the lifetimes obtained for the site-specifically attached 4SA used here were, both for the longest- and for the shortest-lived population, appreciably higher (approximately 5- and 2-fold, respectively) than the respective lifetimes measured for commercially available SA-coated beads (Dynabeads M-270 Streptavidin, Invitrogen/Thermo Fisher)^20^.

This difference may be explained considering that the SA-biotin complex can withstand higher forces when loaded with force from the C-terminus as compared to pulling from the N-terminus, as it has recently been demonstrated for 1SA in AFM SMFS^23^. The attachment of commercially available beads is likely not site-specific, resulting in a variety of pulling geometries, whereas in the custom SA constructs, force was specifically applied from the C-terminus, ensuring highest stability.

## Conclusion

To conclude, we show that the lifetime of the SA-biotin interaction subjected to constant mechanical load strongly depends on the force-loading geometry. Different geometries arise from binding of biotin to one of the four binding pockets of SA and result in lifetimes under force that differ by at least two orders-of-magnitude, despite identical thermodynamic stabilities for binding to the different subunits. Our results illustrate that it is, in general, not possible to infer the mechanical stability of a receptor-ligand complex from its affinity and binding enthalpy.

Such differences between thermal and forced dissociation of molecular complexes are plausible considering the high-dimensional binding energy landscape. Unbinding pathways under mechanical load can be very different from each other and also very different from the thermally preferred ones, as it is also observed e.g. for the force-induced melting of double-stranded DNA in shear- or zipper-geometry^33^. For proteins, similar behavior of monovalent SA has recently been employed by Erlich *et al.* to create a force hierarchy of receptor-ligand complexes^34^. Also, the mechanically most stable receptor-ligand complex measured to date^35^ has just ordinary thermodynamic binding characteristics. Our constant force measurements highlight that there is no general relation between mechanical stability and thermodynamic affinity.

Our work provides a clear route to improving the yield of force spectroscopy experiments and in general of assays where streptavidin-biotin is used for attachment and experiences mechanical loads, e.g. due to fluid flow or magnetic actuation. For measurements utilizing the SA-biotin interaction as a handle, and in particular for constant force SMFS measurements, it is highly beneficial to implement a specific SA tethering geometry that yields long, and single-exponential lifetimes to enable long measurement durations even at high forces. The tethering geometry that we identified as the one yielding the longest lifetimes can be easily realized in experiments by employing the 1SA variant presented in this study. Thus, our results give a straightforward approach for highly specific and stable experiments that employ the streptavidin-biotin linkage.

## Acknowledgements

The authors thank Thomas Nicolaus and Angelika Kardinal for laboratory assistance, Philipp U. Walker for helpful discussions, and Wolfgang Ott, Magnus S. Bauer, and Lukas F. Milles for providing ELP linkers and the ddFLN4 construct, respectively. This project was funded by the Deutsche Forschungsgemeinschaft (DFG, German Research Foundation) Project-ID 201269156, SFB 1032 and Project-ID 386143268, “Unraveling the Mechano-Regulation of Von Willebrand Factor”.

## References

1. O. H. Laitinen, V. P. Hytonen, H. R. Nordlund and M. S. Kulomaa, Cell Mol Life Sci, 2006, 63, 2992–3017.

2. C. M. Dundas, D. Demonte and S. Park, Appl Microbiol Biotechnol, 2013, 97, 9343–9353.

3. M. Wilchek and E. A. Bayer, Anal Biochem, 1988, 171, 1–32.

4. V. V. Didenko, Anal Biochem, 1993, 213, 75–78.

5. F. Kriegel, W. Vanderlinden, T. Nicolaus, A. Kardinal and J. Lipfert, Methods Mol Biol, 2018, 1814, 75–98.

6. I. D. Vilfan, J. Lipfert, D. A. Koster, S. G. Lemay and N. H. Dekker, in Handbook of Single-Molecule Biophysics, eds. P. Hinterdorfer and A. Oijen, Springer US, New York, NY, 2009, DOI: 10.1007/978-0-387-76497-9_13, pp. 371–395.

7. E. A. Bayer and M. Wilchek, Methods Enzymol, 1990, 184, 138–160.

8. M. G. Cull and P. J. Schatz, Methods Enzymol, 2000, 326, 430–440.

9. S. M. Cannizzaro, R. F. Padera, R. Langer, R. A. Rogers, F. E. Black, M. C. Davies, S. J. Tendler and K. M. Shakesheff, Biotechnol Bioeng, 1998, 58, 529–535.

10. V. T. Moy, E. L. Florin and H. E. Gaub, Science, 1994, 266, 257–259.

11. J. Lipfert, J. W. Kerssemakers, T. Jager and N. H. Dekker, Nat Methods, 2010, 7, 977–980.

12. J. Lipfert, M. M. van Oene, M. Lee, F. Pedaci and N. H. Dekker, Chem Rev, 2015, 115, 1449–1474.

13. Jaime A. Rivas-Pardo, Edward C. Eckels, I. Popa, P. Kosuri, Wolfgang A. Linke and Julio M. Fernández, Cell Reports, 2016, 14, 1339–1347.

14. F. Baumann, M. S. Bauer, L. F. Milles, A. Alexandrovich, H. E. Gaub and D. A. Pippig, Nat Nanotechnol, 2016, 11, 89–94.

15. W. Ott, M. A. Jobst, C. Schoeler, H. E. Gaub and M. A. Nash, J Struct Biol, 2017, 197, 3–12.

16. R. Walder, M. A. LeBlanc, W. J. Van Patten, D. T. Edwards, J. A. Greenberg, A. Adhikari, S. R. Okoniewski, R. M. A. Sullan, D. Rabuka, M. C. Sousa and T. T. Perkins, J Am Chem Soc, 2017, 139, 9867–9875.

17. Z. Ganim and M. Rief, Proc Natl Acad Sci U S A, 2017, 114, 11052–11056.

18. M. Krieg, G. Fläschner, D. Alsteens, B. M. Gaub, W. H. Roos, G. J. L. Wuite, H. E. Gaub, C. Gerber, Y. F. Dufrêne and D. J. Müller, Nature Reviews Physics, 2019, 1, 41–57.

19. H. Chen, G. Yuan, R. S. Winardhi, M. Yao, I. Popa, J. M. Fernandez and J. Yan, J Am Chem Soc, 2015, 137, 3540–3546.

20. A. Lof, P. U. Walker, S. M. Sedlak, S. Gruber, T. Obser, M. A. Brehm, M. Benoit and J. Lipfert, Proc Natl Acad Sci U S A, 2019, 116, 18798–18807.

21. S. M. Sedlak, M. S. Bauer, C. Kluger, L. C. Schendel, L. F. Milles, D. A. Pippig and H. E. Gaub, PLOS ONE, 2017, 12, e0188722.

22. G. Fonnum, N. Hofslokken, E. Aksnes, L. Killas, P. Stenstad, R. Schmid, J. Bjorgum, T. Nilsen and A. Berge, EU Patent EP1, 2006, 693.

23. S. M. Sedlak, L. C. Schendel, M. C. R. Melo, D. A. Pippig, Z. Luthey-Schulten, H. E. Gaub and R. C. Bernardi, Nano Letters, 2019, 19, 3415–3421.

24. S. M. Sedlak, L. C. Schendel, H. E. Gaub and R. C. Bernardi, Science Advances, 2020, 6, eaay5999.

25. M. Howarth, D. J. Chinnapen, K. Gerrow, P. C. Dorrestein, M. R. Grandy, N. L. Kelleher, A. El-Husseini and A. Y. Ting, Nat Methods, 2006, 3, 267–273.

26. A. Chilkoti and P. S. Stayton, Journal of the American Chemical Society, 1995, 117, 10622–10628.

27. M. Sarter, D. Niether, B. W. Koenig, W. Lohstroh, M. Zamponi, N. H. Jalarvo, S. Wiegand, J. Fitter and A. M. Stadler, The Journal of Physical Chemistry B, 2019, DOI: 10.1021/acs.jpcb.9b08467.

28. T. R. Strick, J. F. Allemand, D. Bensimon, A. Bensimon and V. Croquette, Science, 1996, 271, 1835–1837.

29. J. Lipfert, X. Hao and N. H. Dekker, Biophys J, 2009, 96, 5040–5049.

30. I. Schwaiger, A. Kardinal, M. Schleicher, A. A. Noegel and M. Rief, Nat Struct Mol Biol, 2004, 11, 81–85.

31. W. Ott, M. A. Jobst, M. S. Bauer, E. Durner, L. F. Milles, M. A. Nash and H. E. Gaub, ACS Nano, 2017, 11, 6346–6354.

32. M. S. Bauer, L. F. Milles, S. M. Sedlak and H. E. Gaub, bioRxiv, 2018.

33. S. K. Kufer, E. M. Puchner, H. Gumpp, T. Liedl and H. E. Gaub, Science, 2008, 319, 594–596.

34. K. R. Erlich, S. M. Sedlak, M. A. Jobst, L. F. Milles and H. E. Gaub, Nanoscale, 2019, 11, 407–411.

35. L. F. Milles, K. Schulten, H. E. Gaub and R. C. Bernardi, Science, 2018, 359, 1527–1533.

36. S. Basu, A. Finke, L. Vera, M. Wang and V. Olieric, Acta Crystallographica Section D, 2019, 75, 262–271.

37. W. Humphrey, A. Dalke and K. Schulten, J Mol Graph, 1996, 14, 33-38, 27-38.

