## Supplementary Information for "Designed Anchoring Geometries Determine Lifetimes of Biotin-Streptavidin Bonds under Constant Load and Enable Ultra-Stable Coupling"

#### Materials and Methods

##### Expression of SA constructs.

Non-functional SA subunits are created by mutating three amino acids (N234A, S27D, S45A) as described by Howarth *et al.*[1] Amino acid sequences of all subunits are provided in the section "Sequences of the protein constructs" below. Functional and non-functional SA subunits were expressed separately and then mixed to obtain SA of different valencies: Tetravalent SA (4SA), trivalent SA (3SA), monovalent SA (1SA), and non-functional SA (0SA). 4SA and 1SA contained a unique cysteine for surface immobilization at the C-terminus of one functional subunit. 3SA and 0SA contained a unique cysteine for surface immobilization at the C-terminus of one non-functional subunit. To select for the correct SA stoichiometry, the cysteine-labeled subunit further contained a polyhistidine tag for purification by nickel-immobilized metal ion affinity chromatography. The unique subunits used for purification and immobilization were incubated either with a 10-fold excess of untagged functional (4SA or 3SA) or untagged non-functional (1SA and 0SA) subunits to assemble the SA tetramers with defined valencies.
All SA subunits were cloned into pET vectors (Merck Millipore, Burlington, USA). SA plasmids were transferred to *E. coli* BL21(DE3)-CodonPlus cells (Agilent Technologies Inc., Santa Clara, USA) and expressed in SB medium. 15 ml of preculture, which was grown overnight at 37°C, was used to inoculate 500 ml of SB medium containing the appropriate antibiotic. Cells were grown at 37°C. At an $OD_{600}$ of 0.8, expression was induced with 0.2 mM IPTG and the temperature was reduced to 18°C for 16 h. The cultures were spun down so that bacterial pellets formed, which were then stored at –80°C.

##### Purification of SA constructs.

All steps except for chromatography were performed on ice or at 4°C, respectively. SA cell pellets were thawed, suspended in 5 ml/g Bacterial Protein Extraction Reagent (B-PER, Thermo Fisher Scientific, Waltham, MA) and incubated with 1 µg lysozyme and 0.05 µg DNAse I per gram bacterial pellet on a rolling shaker for 20 minutes. To ensure full break-up, cells were subsequently sonicated. The lysed cells were then centrifuged. The supernatants were discarded. The pellets were resuspended in lysis buffer (PBS, 1 mM DTT, 0.1% Triton X-100). Sonification, centrifugation and resuspension were repeated four to five times until the supernatant was a clear liquid. The pellets were then resuspended in denaturing buffer (PBS, 6 M guanidine hydrochloride), sonicated and centrifuged. This time, supernatants contained the protein. Supernatants were filtered through a sterile 0.22 µm filter. Then, the absorption at 280 nm was determined. Denatured subunits were mixed in a 1:10 ratio (subunits with and without polyhistidine tag). The mixture was then slowly diluted into 500 ml PBS and stirred overnight using a magnetic stirrer. This solution containing the refolded and reassembled SA was loaded onto a Ni-NTA column (HisTrap FF, GE Healthcare, Chicago, USA). We employed a gradient elution to elute SA from the column and selected for those SA containing a single polyhistidine tag (elution fractions were checked by SDS-PAGE). Elution fractions containing the protein were dialyzed against PBS and then stored in PBS at 4°C. We found SA to be functional (such that successful MT experiments were possible) even after two years of continuous storage in PBS at 4°C after purification.

**Expression and purification of ddFLN4.**

The recombinant ddFLN4 construct[2] expressed in *E.coli* (with the internal cysteine at position 18 mutated to serine) was a kind gift from Lukas F. Milles. At its C-terminus, the ddFLN4 construct possesses a polyhistidine-tag for purification and a ybbR-tag. At its N-terminus, the construct possesses a short linker sequence (MGTGSGSGSGSAGTGSG) with the N-terminal methionine being followed by a single glycine. Due to efficient cleavage of the methionine by *E.coli* methionine aminopeptidases, the glycine is available for Sortase-catalyzed ligation.

The ddFLN4 gene was synthesized codon-optimized for expression in *E.coli* as a linear DNA fragment (GeneArt – Thermo Fisher Scientific, Regensburg, Germany), and inserted into pET28a vectors via Gibson assembly (New England Biolabs, Frankfurt, Germany). Protein expression in *E.coli* NiCo21 (DE3) (New England Biolabs) and purification via the polyhistidine-tag were carried out as previously described[2].

**Preparation of biotinylated DNA.**

To prepare biotinylated DNA, we performed a polymerase chain reaction using a biotinylated primer and a regular DNA reverse primer. As the template DNA, we used the pET28a vector encoding for the functional SA subunit. Primers (BIO-TEG (Triethylenglycol)-5'-atggaagcggggattaccggc-3' and 5'-ctgaccgctccaagtcgtagcg-3') were purchased from Eurofins Genomics, Ebersberg, Germany. For purification of the PCR product, we performed size-exclusion chromatography (with a Superdex 75 Increase 10/300 GL column) on an Äkta Explorer FPLC system (GE Healthcare, Chicago, USA) using PBS as running buffer.

**AFM imaging.**

SA constructs were reduced using 50 mM dithiothreitol and mixed with biotinylated 250 bp double-stranded DNA in PBS buffer, with DNA being in excess to ensure that SA molecules with the maximum possible number of bound DNA strands can be observed. A 1:20 SA:DNA stoichiometry was chosen for 4SA and 3SA, and a 1:4 stoichiometry for 1SA and 0SA, with a final DNA concentration of approximately 4 nM.

Preparation of poly-l-lysine (PLL) coated mica substrates for AFM imaging was performed analogously to a recently described protocol[3, 4]. After at least 1 h of incubation, 20 µl of the SA–DNA mix were incubated on a PLL-coated substrate for 30 s, which was subsequently rinsed with water and finally dried in a gentle stream of nitrogen. The positively charged PLL allows for stable attachment of negatively charged DNA and of DNA-streptavidin complexes. Free streptavidin without bound DNA strands, however, does not stably attach to the substrate. AFM images of 1 µm x 1 µm or 2 µm x 2 µm and 1024 x 1024 pixels were recorded in tapping mode in air, using an MFP-3D AFM (Asylum Research, Santa Barbara, CA) and cantilevers with silicon tips (AC160TS, Olympus, Japan), with a nominal spring constant of 26 N/m and a resonance frequency of approximately 300 kHz. Raw image data were processed using SPIP software (v6.5.1; Image Metrology, Denmark). Image processing involved plane correction (third order polynomial plane-fitting), line-wise flattening (according to the histogram alignment routine), and Gaussian smoothing (for zoom-ins only).

**Isothermal titration calorimetry.**

ITC was performed at 25°C on a MicroCal iTC200 (Malvern, Worcestershire, UK). 8.0 mg Biotin were weighed out and dissolved in 40 ml PBS to obtain a stock solution of about 820 µM.

SA was dissolved in PBS, using Zeba Spin Desalting Columns, 40K MWCO (Thermo Fisher Scientific, Waltham, USA) for buffer exchange. The concentration of SA was determined from the absorption at 280 nm using a spectrophotometer (NanoDrop 1000, Thermo Fisher Scientific, Waltham, USA) and a molar attenuation coefficient of $\varepsilon_{280}$=167,760 $M^{-1}cm^{-1}$. SA was used as analyte, biotin as titrant. A 10-fold concentration of biotin was used for 1SA, a 30-fold excess for 3SA, and a 40-fold excess for 4SA, as the ratio of the measurement cell volume to the total titrant volume is five to one.

**Fitting of the ITC data.**

ITC data are fitted as described in the Appendix of the "ITC Data Analysis in Origin® Tutorial Guide" by MicroCal, LLC. In brief, the fit is approached as follows: First, the concentration of receptor $R_i$ and ligand $L_i$ molecules in the sample cell after each injection $i$ of volume $\Delta V_i$ has to be calculated. With $V_0$ as the volume of the measurement cell and $X$ as the ligand concentration in the syringe, we can set up the following relations

$$R_{i+1} \cdot V_0 = R_i \cdot V_0 - 0.5 \cdot (R_{i+1} + R_i) \cdot \Delta V_i$$

$$L_{i+1} \cdot V_0 = L_i \cdot V_0 - 0.5 \cdot (L_{i+1} + L_i) \cdot \Delta V_i + X \cdot \Delta V_i$$

These formulas take into account that with each injection of volume $\Delta V_i$, the same volume $\Delta V_i$ is displaced from the measurement cell containing both ligands and receptor at the current concentration – on average $0.5 \cdot (R_{i+1} + R_i)$.

We obtain the recursive relations

$$R_{i+1} = R_i \cdot \frac{V_0 - 0.5\ \cdot \Delta V_i}{V_0 + 0.5\ \cdot \Delta V_i}$$

$$L_{i+1} = \frac{X \cdot \Delta V_i + L_i \cdot (V_0 - 0.5\ \cdot \Delta V_i)}{V_0 + 0.5\ \cdot \Delta V_i}$$

For a set of identical binding sites, the fraction bound for each $R_i$ and $L_i$, can be calculated according to

$$f = 0.5 \cdot \left(1 + \frac{L_i}{n \cdot R_i} + \frac{K_D}{n \cdot R_i} - \sqrt{\left(1 + \frac{L_i}{n \cdot R_i} + \frac{K_D}{n \cdot R_i}\right)^2 - \frac{4 \cdot L_i}{n \cdot R_i}}\right)$$

where $n$ is the stoichiometry and $K_D$ is the affinity constant.

The total heat $Q$ that would be released going from $(R_0, L = 0)$ to $(R_i, L_i)$ is then calculated by

$$Q = n \cdot f \cdot H \cdot R_i \cdot V_0$$

where $H$ is the binding enthalpy per mole.

In the experiment not the total heat $Q$, but the difference in heat between two injections is measured

$$\Delta Q_i = Q_i - Q_{i-1} + 0.5 \cdot (Q_{i-1} + Q_i) \cdot \frac{\Delta V_i}{V_0}$$

where the last term takes into account that also the molecules contained in the replaced volume $\Delta V_i$ contribute to the heat while being pushed out of the measurement cell.

For fitting the data, initial values for $n$, $K_D$ and $H$ are guesses. Then all $\Delta Q_i$ for all injections are calculated and compared with the experimental values. The values for $n$, $K_D$ and $H$ are then improved and the procedure is repeated until no further improvement of the fit occurs.

We note that due to the discretization of the measurement, the fit does not represent an ideal sigmoidal binding curve (which would be obtained for an infinite number of infinitesimal injections). Instead, the data points represent discrete heat differences between distinct injections, *i.e.* they can be rather seen as an average over a certain part of the ideal binding curve (**Supplementary Fig. S7**). Due to the high affinity of the SA-biotin interaction, the saturation of receptors is very abrupt and leads to a sudden change in the released heat. Averaging over this part of the ideal binding curve leads to data points that are no longer on the ideal binding curve. Since the heat released in a single injection is always plotted against the final molecular ratio (the ratio after the injection), this discretized curve is shifted to the right of the underlying ideal binding curve (**Supplementary Fig. S7**). Nevertheless, $n$ and $H$ can be reliably obtained from the fit, which takes into account the discretization. The true value for $K_D$ can however (for the SA-biotin system) not be measured.

**Functionalization of magnetic beads with SA constructs.**
5 µM of 1SA, 3SA, or 4SA were supplemented with 5 mM Bond-Breaker TCEP Solution (Thermo Fisher Scientific). After 30 minutes, the mixture was purified using Zeba Spin Desalting Columns, 40K MWCO (Thermo Fisher Scientific) equilibrated with coupling buffer (50 mM NaCl, 50 mM $NaHPO_4$, 10 mM EDTA, pH 7.2) according to the manufacturer's instructions.
Bifunctional polyethylene glycol of 5,000 g/mol molecular weight with an N-hydroxysuccinimide group at one end and a maleimide group at the other (NHS-PEG5000-MAL, Rapp Polymere, Tübingen, Germany) was dissolved in 50 mM HEPES, pH 7.5, to a final concentration of 25 mM and immediately used to incubate superparamagnetic beads with amine groups (Dynabeads M-270 Amine, Invitrogen/Thermo Fisher). After 45 min, beads were washed extensively first with DMSO and then with ultrapure water. Beads were then incubated with the respective SA construct in coupling buffer for 90 min and extensively washed with measurement buffer (20 mM HEPES, 150 mM NaCl, 1 mM $MgCl_2$, 1 mM $CaCl_2$, 0.1% (v/v) Tween-20, pH 7.4).

**Magnetic tweezers setup.**
MT experiments were performed on a previously described custom setup[5, 6]. The setup employs a pair of permanent magnets (5×5×5 $mm^3$ each; W-05-N50-G, Supermagnete, Switzerland) in vertical configuration[7]. The distance between magnets and flow cell (and, thus, the force) is controlled by a DC-motor (M-126.PD2; PI Physikinstrumente, Germany). An LED (69647, Lumitronix LED Technik GmbH, Germany) is used for illumination. A 40x oil immersion objective (UPLFLN 40x, Olympus, Japan) and a CMOS sensor camera with 4096×3072 pixels (12M Falcon2, Teledyne Dalsa, Canada) allow to image a large field of view of approximately 440 × 330 $\mu m^2$ at a frame rate of 58 Hz. Images are transferred to a frame grabber (PCIe 1433; National Instruments, Austin, TX) and analyzed with a LabView-based open-source tracking software[8]. The bead tracking accuracy of the setup is ≈0.6 nm in ($x$, $y$) and ≈1.5 nm in $z$ direction. For creating the look-up table required for tracking the bead positions in $z$, the objective is mounted on a piezo stage (Pifoc P-726.1CD, PI Physikinstrumente). Force calibration was conducted as described by te Velthuis *et al.*[9] based on the transverse fluctuations of long DNA tethers. Importantly, for the small extension changes on the length scales of our protein tethers, the force stays essentially constant[6], with the relative change in force due to tether stretching or protein unfolding being $< 10^{-4}$. Force deviations due to magnetic field inhomogeneities across the full range of the field of view are < 3%. The largest source of force

uncertainty is the bead-to-bead variation, which is on the order of ≤ 10% for the beads used in this study[6, 7, 10, 11].

**Magnetic Tweezers experiments.**

Preparation of flow cells was performed as described[6]. In brief, aminosilanized glass slides were functionalized with elastin-like polypeptide (ELP) linkers[12], possessing a single cysteine at their N terminus as well as a C-terminal Sortase motif, via a small-molecule crosslinker with a thiol-reactive maleimide group [sulfosuccinimidyl 4-(N-maleimidomethyl)cyclohexane-1-carboxylate; Sulfo-SMCC, Thermo Fisher Scientific]. Flow cells were then assembled from an ELP-functionalized slide as bottom and a non-functionalized glass slide with two small holes for inlet and outlet as top, with a layer of cut-out parafilm (Pechiney Plastic Packaging Inc., Chicago, IL) in between to form a channel. Flow cells were incubated with 1% casein solution (Sigma-Aldrich) for 1 h and flushed with 1 ml (approximately 20 flowcell volumes) of buffer (20 mM HEPES, 150 mM NaCl, 1 mM $MgCl_2$, 1 mM $CaCl_2$, pH 7.4).

CoA-biotin (New England Biolabs) was coupled to the ybbR-tag of the ddFLN4 construct in a bulk reaction in the presence of 5 μM sfp phosphopantetheinyl transferase and 10 mM $MgCl_2$ at 37 °C for 60 min. Afterwards, ddFLN4 was diluted to a final concentration of approximately 20 nM in 20 mM HEPES, 150 mM NaCl, 1 mM $MgCl_2$, 1 mM $CaCl_2$, pH 7.4, and incubated in the flow cell in the presence of 2 μM evolved pentamutant Sortase A[13, 14] for 30 min. Subsequently, the flow cell was flushed with 1 ml of measurement buffer (20 mM HEPES, 150 mM NaCl, 1 mM $MgCl_2$, 1 mM $CaCl_2$, 0.1% (v/v) Tween-20, pH 7.4). Finally, beads functionalized with the respective SA construct were incubated in the flow cell for 60 s, and unbound beads were flushed out with 2 ml of measurement buffer.

At the beginning of each measurement, the tethered beads were subjected to two 5-min intervals of a constant force of 25 pN to allow for identification of specific, single-tethered beads by the characteristic two-step unfolding pattern of ddFLN4[6, 15] (**Fig. 3**). Only beads that showed the ddFLN4 pattern were analyzed further. Importantly, essentially no specifically attached beads ruptured during this phase of the measurement. After 30 s at a low resting force of 0.5 pN, beads were subjected to a constant force of 65 pN for either 15 h (4SA and 1SA) or 5 h (3SA), and the time until bead rupture was monitored. All measurements were performed at room temperature (≈ 22°C).

**Analysis of Magnetic Tweezers measurements.**

Lifetimes were determined from the survival fraction *vs.* time data based on > 50 tether rupture events for each SA construct. To estimate the robustness of fits and determine error bounds for the fitted parameters, we used a bootstrapping procedure. From each experimental data set, 1000 synthetic data sets of the same size were generated by random drawing of data points from the experimental data set with repeats. Synthetic data sets were fit individually and the uncertainties on the final parameters reported are standard deviations over the bootstrap ensemble.

We tested different models for each data set. The models, fitted parameters, and quality of fits are summarized in the table below. The quality of the fits was evaluated by considering the residuals defined as

$$Res = \frac{\left(\sum_{i=1}^{N}\left(f_{exp,i}(t_i) - f_i(t_i)\right)^2\right)^{1/2}}{N}$$

where $f_{exp,i}$ are the measured survival fractions and $f_i$ are the modeled survival fraction at times $t_i$. $N$ is the number of data points in each data set. The models used in **Fig. 3** in the main text are highlighted by the dark shading in the table below.

| **Data set** | **Model** | **Fitting Parameter** | **Resi-duals** |
|---|---|---|---|
| 1SA | $f(t) = f_1 \cdot \exp(-t/\tau_1)$ | $f_1 = 0.92 \pm 0.01$<br>$\tau_1 = 2.61 \cdot 10^4$ s $\pm$ 680 s | 0.0041 |
| | $f(t) = \exp(-t/\tau_1)$ | $\tau_1 = 2.25 \cdot 10^4$ s | 0.0073 |
| 3SA | $f(t) = f_2/3\,[\exp(-t/\tau_2) + 2\cdot\exp(-t/\tau_3)]$ | $f_2 = 0.95 \pm 0.01$<br>$\tau_2 = 3.61 \cdot 10^3$ s $\pm$ 350 s<br>$\tau_3 = 199$ s $\pm$ 10 s | 0.0034 |
| | $f(t) = 1/3\,[\exp(-t/\tau_2) + 2\cdot\exp(-t/\tau_3)]$ | $\tau_2 = 3.36 \cdot 10^3$ s<br>$\tau_3 = 167$ s | 0.0042 |
| | $f(t) = f_2/3\,[\exp(-t/\tau_A) + \exp(-t/\tau_B) + \exp(-t/\tau_C)]$ | $f_2 = 0.97 \pm 0.01$<br>$\tau_A = 98$ s $\pm$ 12 s<br>$\tau_B = 365$ s $\pm$ 30 s<br>$\tau_C = 3.1 \cdot 10^3$ s $\pm$ 230 s | 0.0027 |
| | $f(t) = f_2 \cdot \exp(-t/\tau_2)$ | $f_2 = 0.87$<br>$\tau_2 = 560$ s | 0.010 |
| 4SA | $f(t) = f_3/4[\exp(-t/\tau_1) + \exp(-t/\tau_2) + 2\cdot\exp(-t/\tau_3)]$ | $f_3 = 0.86 \pm 0.01$ | 0.0073 |
| | $f(t) = 1/4\,[\exp(-t/\tau_1) + \exp(-t/\tau_2) + 2\cdot\exp(-t/\tau_3)]$ | -None- | 0.0142 |

### Sequences of the protein constructs

**Functional SA subunit:**

MEAGITGTWYNQLGSTFIVTAGADGALTGTYESAVGNAESRYVLTGRYDSAPATDG
SGTALGWTVAWKNNYRNAHSATTWSGQYVGGAEARINTQWLLTSGTTEANAWKS
TLVGHDTFTKVKPSAAS

**Functional SA subunit with C-terminal cysteine and His-tag:**

MEAGITGTWYNQLGSTFIVTAGADGALTGTYESAVGNAESRYVLTGRYDSAPATDG
SGTALGWTVAWKNNYRNAHSATTWSGQYVGGAEARINTQWLLTSGTTEANAWKS
TLVGHDTFTKVKPSAASCLEHHHHHH

**Non-functional SA subunit:**

MEAGITGTWYAQLGDTFIVTAGADGALTGTYEAAVGNAESRYVLTGRYDSAPATDG
SGTALGWTVAWKNNYRNAHSATTWSGQYVGGAEARINTQWLLTSGTTEANAWKS
TLVGHDTFTKVKPSAAS

**Non-functional SA subunit with C-terminal cysteine and His-tag:**

MEAGITGTWYAQLGDTFIVTAGADGALTGTYEAAVGNAESRYVLTGRYDSAPATDG
SGTALGWTVAWKNNYRNAHSATTWSGQYVGGAEARINTQWLLTSGTTEANAWKS
TLVGHDTFTKVKPSAASCLEHHHHHH

**ddFLN4 (C18S) construct with N-terminal glycine and short linker sequence, and C-terminal His-tag and ybbR-tag:**

MGTGSGSGSGSAGTGSGADPEKSYAEGPGLDGGESFQPSKFKIHAVDPDGVHRTDG
GDGFVV
TIEGPAPVDPVMVDNGDGTYDVEFEPKEAGDYVINLTLDGDNVNGFPKTVTVKPAPS
G HHHHHHGSDSLEFIASKLA

### Supplementary Figures

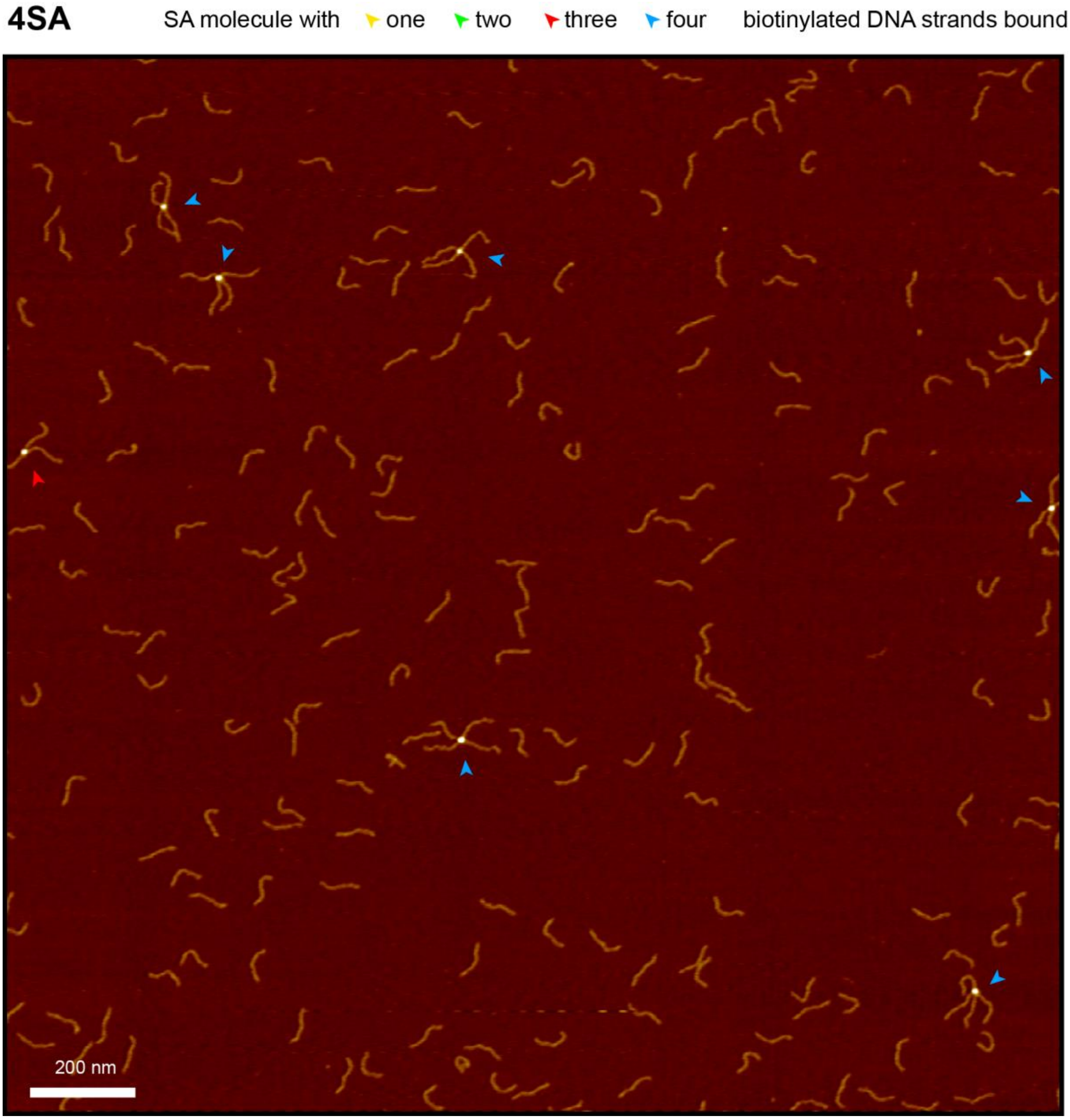


**Supplementary Fig. S1. AFM imaging of 4SA with biotinylated DNA.**
Representative AFM image of 4SA and biotinylated 250 bp dsDNA after incubation in a 1:20 ratio. Arrowheads mark streptavidin molecules, with the color of arrowheads indicating the number of bound DNA strands (yellow – one, green – two, red – three, blue – four). For 4SA, up to four bound strands were observed. Height range of color scale is 2.5 nm.

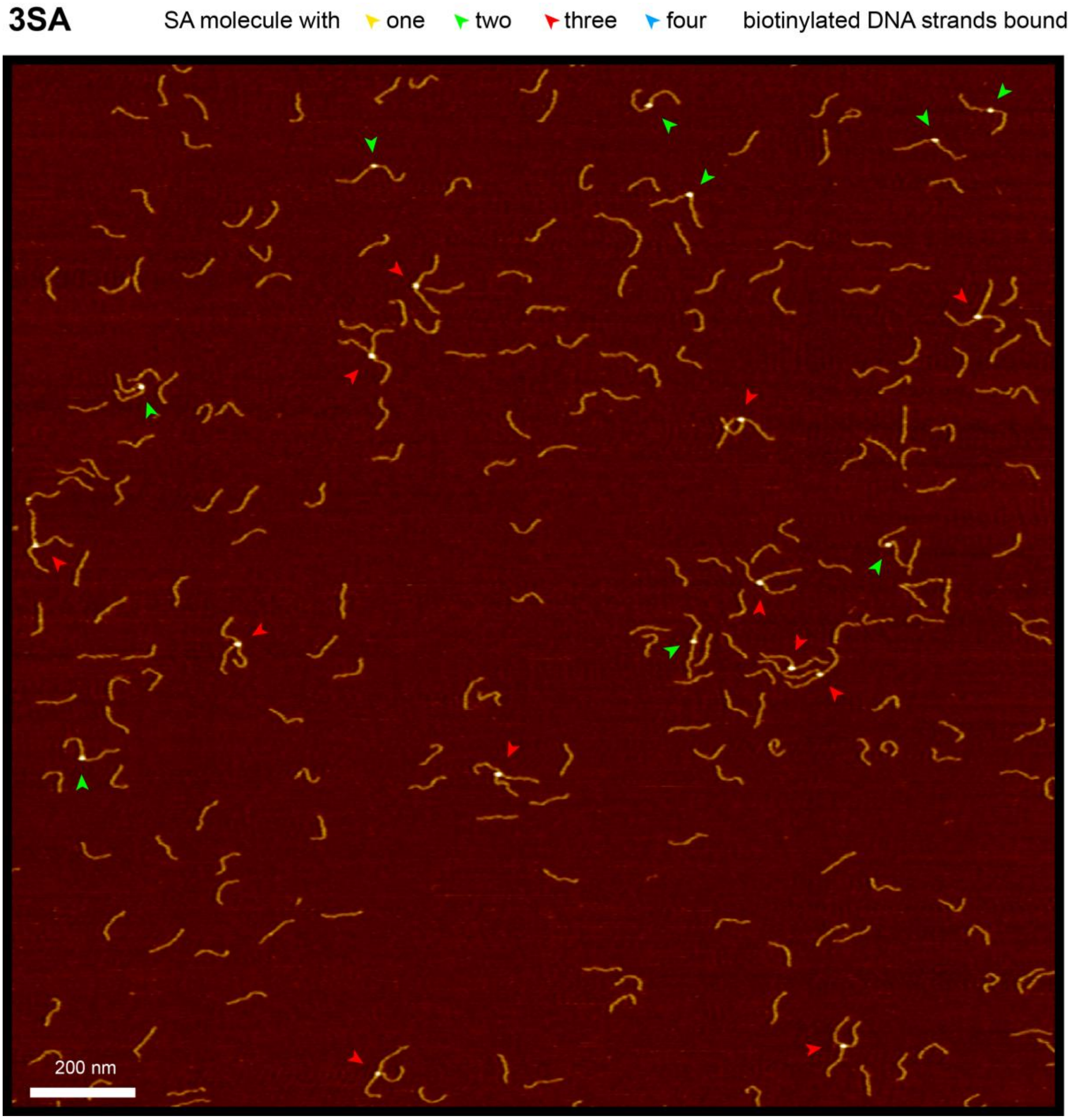


**Supplementary Fig. S2. AFM imaging of 3SA with biotinylated DNA.**
Representative AFM image of 3SA and biotinylated 250 bp dsDNA after incubation in a 1:20 ratio. Arrowheads mark streptavidin molecules, with the color of arrowheads indicating the number of bound DNA strands (yellow – one, green – two, red – three, blue – four). For 3SA, up to three bound strands were observed. No 3SA molecules with four strands were observed. Height range of color scale is 2 nm.

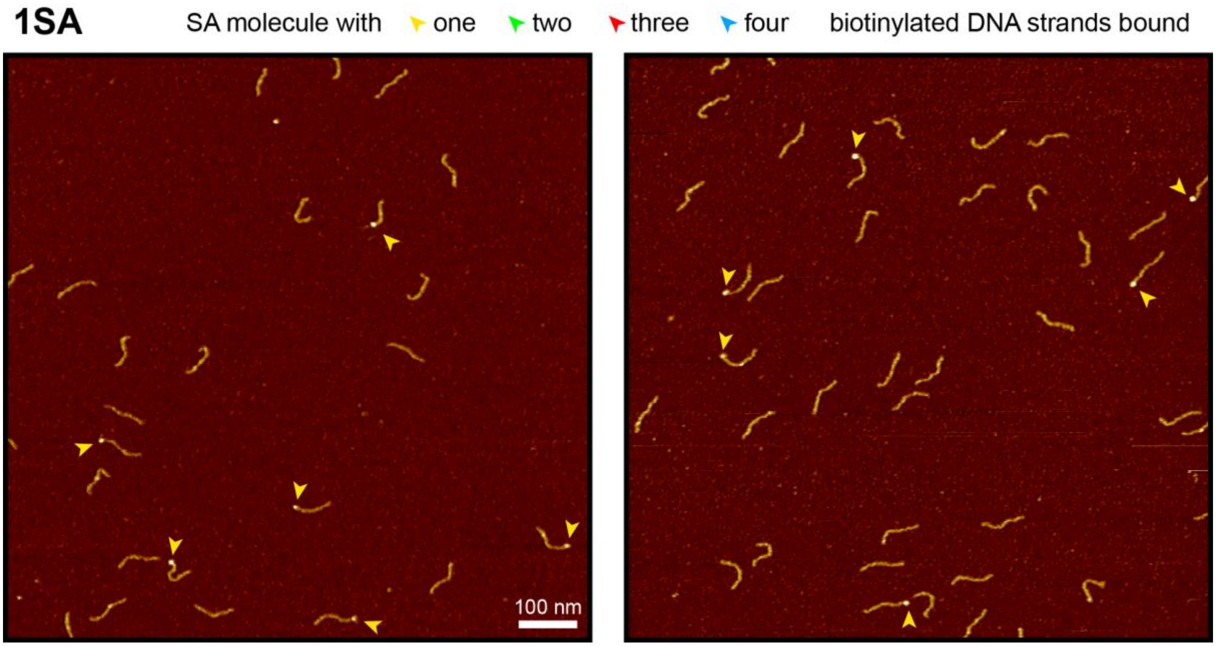


**Supplementary Fig. S3. AFM imaging of 1SA with biotinylated DNA.**
Representative AFM images of 1SA and biotinylated 250 bp dsDNA after incubation in a 1:4 ratio. Arrowheads mark streptavidin molecules, with the color of arrowheads indicating the number of bound DNA strands (yellow – one, green – two, red – three, blue – four). Only 1SA molecules with a single bound DNA strand were observed. Height range of color scale is 2 nm.

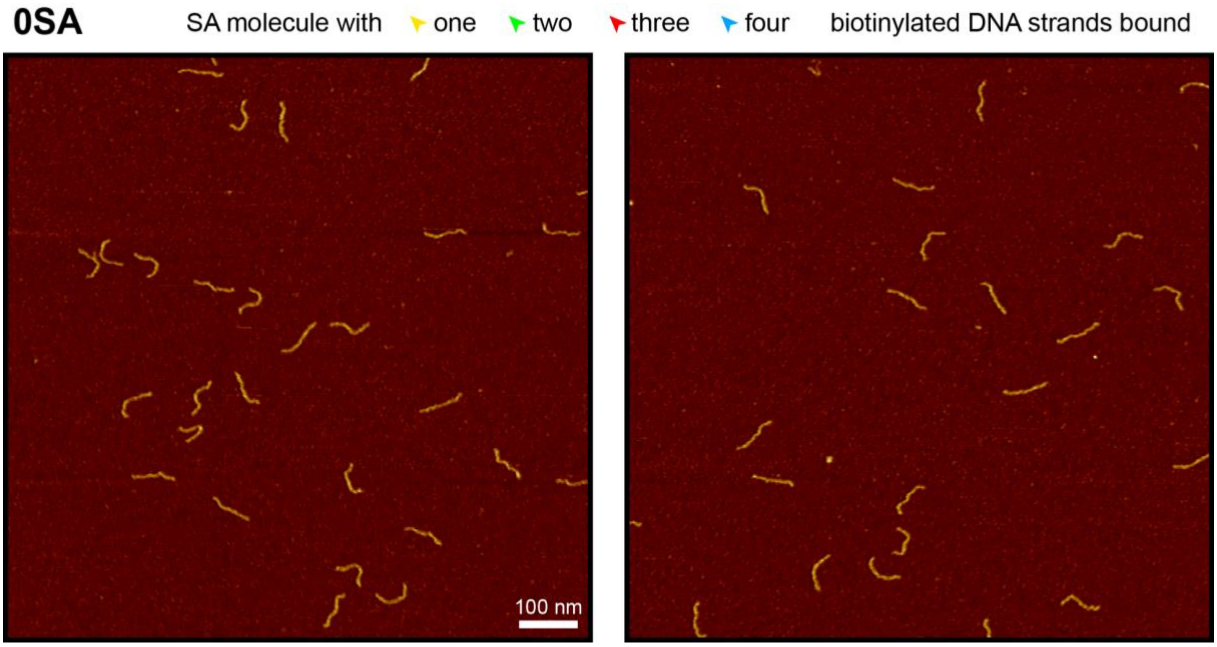


**Supplementary Fig. S4. AFM imaging of 0SA with biotinylated DNA.**
Representative AFM images of 0SA and biotinylated 250 bp dsDNA after incubation in a 1:4 ratio. Only free DNA strands not bound to 0SA were observed. Free 0SA molecules could not be observed since they do not stably attach to the positively charged poly-L-lysine coated mica substrate. Height range of color scale is 2 nm.

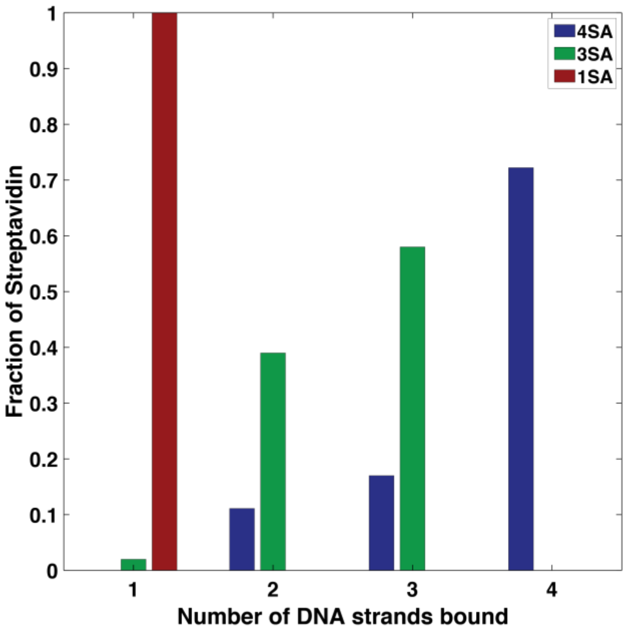


**Supplementary Fig. S5. Number of biotinylated DNA strands bound to SA of different valencies.** Numbers obtained from AFM images as described in **Supplementary Materials and Methods**. Total number of SA molecules observed: 1SA: 33, 3SA: 43, 4SA: 18. AFM images confirm the valency of the SA molecules: The number of biotinylated DNA strands bound to one SA molecule is strictly limited by the number of functional subunits.

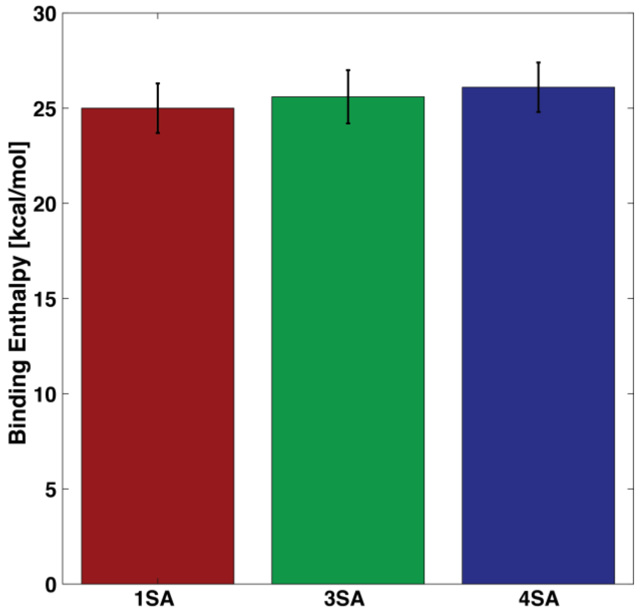


**Supplementary Fig. S6. Binding enthalpy per binding site of SA of different valencies.** Binding enthalpies per binding site for biotin binding to different SA variants, determined from fits to the ITC data (Fig. 2c). Error bars take into account the uncertainty in the stock concentrations, assuming an uncertainty of 10% in SA concentration and a 5% uncertainty in biotin concentration. Within experimental errors, the binding enthalpies for all SA variants are the same, suggesting that in the absence of force all subunits are equivalent with regard to biotin binding and that there are no effects of binding geometry or binding cooperativity.

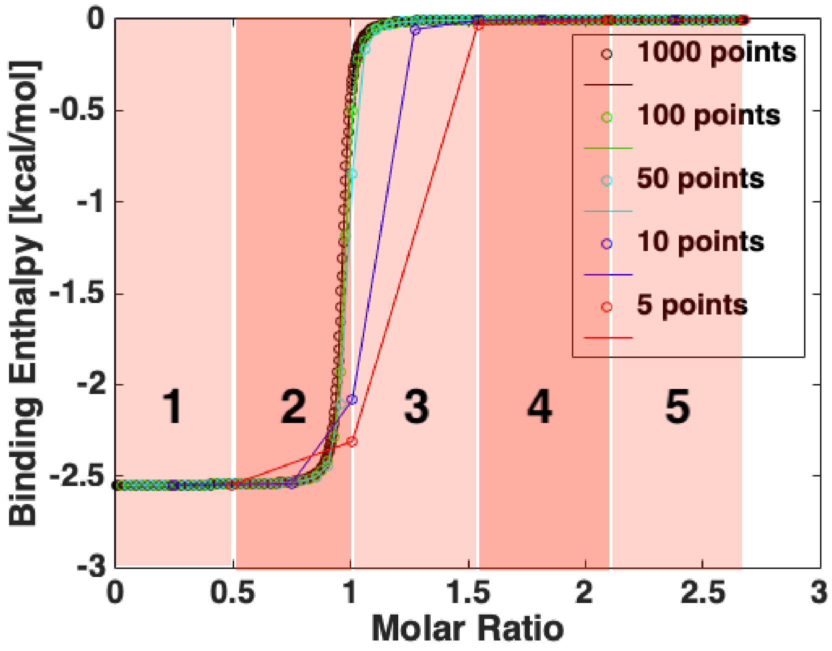


**Supplementary Fig. S7. Discretization effects in ITC binding curves.**

Theoretical binding curve for $n = 1.0$, $K_D = 10^{-9}$ M and $H = -2.5$ kcal/mol. For a large number of very small injections, an ideal binding curve is obtained (black). With small numbers of injections (blue or red), the curve deviates from the ideal binding curve. For only five injections (red), different parts of the curve, which are averaged, are indicated by the numbered red areas in the background of the plot. For flat parts of the curve (area 1, 4 and 5) the discretization only has minor effects, while for the steep parts of the curve (area 2 and 3) the average points are no longer on the ideal curve. Fitting the discretized data, $H$ and $n$ can still be reliably determined. However, the value for the affinity $K_D$ can not be reliably obtained. See the "Fitting of the ITC data" section above for details.

### References


1. M. Howarth, D. J. Chinnapen, K. Gerrow, P. C. Dorrestein, M. R. Grandy, N. L. Kelleher, A. El-Husseini and A. Y. Ting, *Nat Methods*, 2006, **3**, 267-273.
2. L. F. Milles, E. A. Bayer, M. A. Nash and H. E. Gaub, *J Phys Chem B*, 2016, DOI: 10.1021/acs.jpcb.6b09593.
3. J. P. Muller, A. Lof, S. Mielke, T. Obser, L. K. Bruetzel, W. Vanderlinden, J. Lipfert, R. Schneppenheim and M. Benoit, *Biophys J*, 2016, **111**, 312-322.
4. T. Brouns, H. De Keersmaecker, S. F. Konrad, N. Kodera, T. Ando, J. Lipfert, S. De Feyter and W. Vanderlinden, *ACS Nano*, 2018, **12**, 11907-11916.
5. P. U. Walker, W. Vanderlinden and J. Lipfert, *Physical Review E*, 2018, **98**, 042412.
6. A. Lof, P. U. Walker, S. M. Sedlak, S. Gruber, T. Obser, M. A. Brehm, M. Benoit and J. Lipfert, *Proc Natl Acad Sci U S A*, 2019, **116**, 18798-18807.
7. J. Lipfert, X. Hao and N. H. Dekker, *Biophys J*, 2009, **96**, 5040-5049.
8. J. P. Cnossen, D. Dulin and N. H. Dekker, *Rev Sci Instrum*, 2014, **85**, 103712.
9. A. J. W. Te Velthuis, J. W. J. Kerssemakers, J. Lipfert and N. H. Dekker, *Biophysical journal*, 2010, **99**, 1292-1302.
10. I. De Vlaminck, T. Henighan, M. T. van Loenhout, D. R. Burnham and C. Dekker, *PLoS One*, 2012, **7**, e41432.
11. E. Ostrofet, F. S. Papini and D. Dulin, *Sci Rep*, 2018, **8**, 15920.
12. W. Ott, M. A. Jobst, M. S. Bauer, E. Durner, L. F. Milles, M. A. Nash and H. E. Gaub, *ACS Nano*, 2017, **11**, 6346-6354.
13. I. Chen, B. M. Dorr and D. R. Liu, *Proc Natl Acad Sci U S A*, 2011, **108**, 11399-11404.
14. E. Durner, W. Ott, M. A. Nash and H. E. Gaub, *ACS Omega*, 2017, **2**, 3064-3069.
15. I. Schwaiger, A. Kardinal, M. Schleicher, A. A. Noegel and M. Rief, *Nat Struct Mol Biol*, 2004, **11**, 81-85.
